# A DERIVED RELAXATION CONTRAST FROM SYNTHETIC MRI FOR DETECTING NETWORK MICROSTRUCTURAL VULNERABILITY

**DOI:** 10.64898/2026.04.08.717271

**Authors:** Anupa Ekanayake, Scott Hwang, Senal Peiris, Rommy Elyan, Mustafa Ghulam, Mark Tulchinsky, Jianli Wang, Paul Eslinger, Qing Yang, Prasanna Karunanayaka

**Affiliations:** Department of Radiology, Pennsylvania State University College of Medicine, Hershey, PA, USA; Department of Neurology, Pennsylvania State University College of Medicine, Hershey, PA, USA; Department of Neurosurgery, Pennsylvania State University College of Medicine, Hershey, PA, USA

**Keywords:** Myelin, FLAIR MRI, Double Inversion Recovery, MCI, Olfaction

## Abstract

**Background:** Odor identification impairment is an early marker of Alzheimer’s disease (AD) that predicts memory decline, yet its underlying microstructural basis remains unclear. We hypothesized that mild cognitive impairment (MCI) involves early myelin and lipid disruption within olfactory–limbic circuits, detectable using a synthetic MRI–derived contrast that provides complementary sensitivity to myelin volume fraction (MVF).

**Methods:** Thirty-three older adults (healthy controls [HC], n = 16; mild cognitive impairment [MCI], n = 17) completed olfactory and cognitive testing and underwent 3T brain MRI using a QALAS sequence. An MVF map and synthetic FLAIR and DIR images were generated, and a FLAIR–DIR-derived metric (FD) was computed as FD = (FLAIR − DIR) / FLAIR. We investigated ROI-based group differences in olfactory–limbic gray-matter regions and associated white-matter tracts, voxel-wise regressions investigating FD–odor identification associations, and ROI-based MCI vs HC classification using cross-validated logistic regression models.

**Results:** Compared with HC, MCI showed significantly lower FD across olfactory–limbic gray-matter regions and white-matter pathways—including hippocampus, amygdala, orbitofrontal cortex, thalamus, and corpus callosum—whereas MVF differences were more limited. FD achieved moderate discrimination, with baseline performance comparable to MVF. Voxel-wise analyses revealed that better odor identification was associated with higher FD in the hippocampus/parahippocampal and insula; the association persisted after adjusting for voxel-wise MVF. MVF also showed significant positive voxel-wise associations with odor identification in the insula and genu of the corpus callosum.

**Conclusion:** FD is a practical, myelin- and lipid-sensitive contrast derived from routinely acquired synthetic FLAIR & DIR images that complement quantitative MVF. It captures behaviorally relevant variance beyond local myelin content and may improve detection of early olfactory–limbic microstructural changes in MCI. These findings support FD as a scalable candidate marker linking early network disruption to olfactory symptoms across the AD continuum.

## INTRODUCTION

Myelin degeneration is increasingly recognized as a main feature of Alzheimer’s disease (AD) pathology, including during the prodromal stage of mild cognitive impairment (MCI) [1–8]. Multi-parametric mapping via SyMRI enables estimation of myelin content using the myelin volume fraction (MVF), a validated proxy that correlates with histological measures and other myelin metrics [9–11]. However, acquiring the specialized sequences required for MVF mapping is not always feasible in a routine clinical MRI protocol.

Deriving surrogate myelin-sensitive measures from standard clinical MRI contrasts, therefore, holds substantial scientific and translational value [2]. In this context, the ratio of T1- and T2-weighted images (T1w/T2w) has been used to approximate myelin content and has demonstrated strong correlation with quantitative MVF estimates (Spearman ρ ≈ 0.89) [2,10].

We introduce FD, a semi-quantitative MRI contrast metric that leverages the complementary sensitivity of synthetic FLAIR and DIR images to generate a novel tissue characterization index. DIR applies two inversion pulses to suppress both CSF and normal-appearing white matter, thereby accentuating cortical and juxtacortical signal differences [12,13]. FLAIR suppresses cerebrospinal fluid (CSF) signal while providing strong T2-weighting, improving lesion conspicuity and enhancing white–gray matter contrast near CSF interfaces [14]. By using these two contrasts, FD is computed as a normalized difference, FD = (FLAIR – DIR) / FLAIR, yielding a voxel-wise map in which higher values indicate areas where FLAIR signal exceeds DIR signal. This contrast highlights tissue where preserved FLAIR signal that is suppressed on DIR—effectively highlighting myelin-/lipid-rich areas. Since both the FLAIR and DIR contrasts are governed by T1 and T2 relaxation processes, FD effectively captures differences in inversion-recovery signal behavior that may reflect underlying tissue microstructure.

Like the T_1_/T_2_ ratio used in quantitative MRI, FD is sensitive to alterations in myelin, lipid composition, and water content, providing a practical marker of microstructural disruption derived from routinely generated contrasts [15–17]. FD provides a myelin-/ lipid-sensitive contrast across both the white and gray matter. FD likely reflects combined contributions from myelin, lipid membranes, water content, and other tissue properties rather than absolute quantification of myelin. Consistent with this interpretation, FD is expected to demonstrate partial spatial overlap with MVF while also capturing complementary variance not fully reflected by MVF.

As a practical application of FD, we focused on olfactory identification deficits, an established early marker of AD and its MCI phase [18–20]. Impaired odor identification is associated with atrophy in olfactory and limbic regions, including the olfactory cortex, hippocampus, amygdala, entorhinal cortex, and insula [21–23]. Postmortem AD studies have further demonstrated substantial loss of myelinated axons in the olfactory tract [24]. Clinically, poorer odor identification predicts progression from MCI to AD dementia [20,25]. Together, these findings position olfactory performance as a sensitive proxy of regional structural and microstructural integrity, highlighting a biologically grounded context in which FD may offer translational utility [21].

Here, we compare FD with MVF in an MCI cohort, focusing on olfactory and limbic regions. Our goals are to: (1) examine group differences in FD and MVF between healthy controls (HC) and MCI; (2) evaluate classification performance of ROI-based FD versus MVF for distinguishing MCI from HC; (3) investigate voxel-wise associations between FD and olfactory identification scores; and (4) assess the correspondence between FD and MVF across the brain. We hypothesize that FD will detect MCI-related abnormalities in myelin-rich olfactory and limbic regions, correlate with behavioral measures such as odor identification (odor ID), and show spatial overlap with MVF while offering complementary microstructural information not fully captured by MVF.

## METHODS

### Physics of FD

Both the DIR and FLAIR are inversion-recovery–based contrasts. Their signals can be approximated as 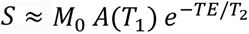, where *A*(*T*_1_)depends on the inversion timings and TR. Under standard assumptions (perfect inversions and spoiled readout), 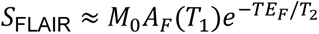 and 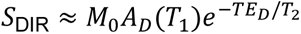. Substituting into *FD* = (FLAIR − DIR)/FLAIR yields 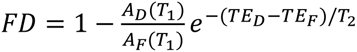, such that proton density (*M*_0_) cancels, and FD emphasizes differences in inversion-recovery weighting (and any TE mismatch) between the two contrasts. FD indirectly depends on T1 and T2 relaxation properties, because both DIR and FLAIR signals are governed by T1 and T2. As mentioned in the introduction, FD reflects differences in inversion-recovery signal behavior linked to tissue microstructure. Analogous to the T1/T2 ratio [2,10,26], it is sensitive to changes in myelin integrity, lipid composition, and water content, thereby serving as a scalable marker of microstructural disruption (see the supplementary material for details).

### Participants

Participants were recruited through Penn State Healthy-affiliated memory clinics and community outreach initiatives. All participants provided written informed consent, and the study protocol was approved by the Penn State College of Medicine Institutional Review Board (IRB). A total of 33 older adults were included (HC = 16, MCI =17) in the study (Table 1). HC met the inclusion criteria of no cognitive complaints, a Mini-Mental State Examination (MMSE) score between 24–30, and a Clinical Dementia Rating (CDR) = 0 with Memory Box=0. MCI participants met consensus clinical criteria, including memory-cognitive concerns reported by participants, an informant, or a clinician, MMSE =18–23), and CDR=0.5 with Memory Box ≥0.5.

**Table 1.** Demographic and clinical characteristics of the healthy control (HC) and mild cognitive impairment (MCI) groups.

|  | HC (n=16) | MCI (n=17) | P |
| --- | --- | --- | --- |
| Gender (M/F) | (3/13) | (10/7) | 0.0185 |
| Age (year) | 63.50 ± 6.55 | 70.35 ± 8.09 | 0.0118 |
| Educational | 17.38 ± 2.12 | 16.62 ± 3.26 | 0.4325 |
| Odor Identification | 16.50 ± 2.16 | 13.94 ± 4.49 | 0.0465 |

Exclusion criteria included limited English proficiency, current smoking, or Geriatric Depression Scale score greater than 6, neurological disorders unrelated to suspected AD, including Parkinson’s disease, multiple sclerosis, Huntington’s disease, or traumatic brain injury with lasting deficits; MRI contraindications (metal implants, pacemaker, or claustrophobia); major psychiatric disorders (schizophrenia, bipolar disorder, or major depression); psychotropic medication use within the past 3 months; substance/alcohol dependence within the past 2 years; or neurodevelopmental disorders (autism spectrum disorder or ADHD).

### Behavioral Testing

Odor identification (ID) was assessed using the computerized OLFACT™ Test Battery (Osmic Enterprises, Inc., Cincinnati, OH), employing a four-alternative forced-choice design. Odor stimuli were delivered using an ETT olfactometer (Emerging Tech Trans, LLC, College Station, TX, USA). Participants identified 20 distinct odors, selecting the correct response from four options at each trial; the total number correct was used as the odor identification score.

### MRI Acquisition

#### T1-weighted Structural Imaging

High-resolution anatomical images were acquired using a T1-weighted MPRAGE sequence for cortical segmentation and anatomical registration. The key acquisition parameters were repetition time (TR) = 1540 ms, echo time (TE) = 2.34 ms, inversion time (TI) = 807 ms, and flip angle = 9°. Images were acquired with 1.0 mm³ isotropic resolution across 176 slices, with a field of view (FOV) of 256 mm and an acceleration factor of 2 using GRAPPA parallel imaging. The total acquisition time was approximately 3.5 minutes.

#### Quantitative Myelin Imaging

Quantitative myelin imaging was performed using a synthetic MRI (SyMRI) protocol implemented via a QALAS-based multi-parametric acquisition [27]. SyMRI models each voxel as a combination of four tissue compartments, namely, myelin (MyC), cellular tissue, excess parenchymal water, and free water, based on their relaxation properties (R1, R2, and proton density). This approach allows for direct voxel-wise estimation of MVF content with high reproducibility and biological specificity.

The acquisition was based on the QALAS sequence, which is a rapid multi-delay, multi-flip angle sequence designed to simultaneously estimate T1, T2, and proton density (PD) in a single acquisition. The key parameters for QALAS acquisition included TR = 4500 ms, TE = 2.24 ms, and flip angles of 4° across five inversion times (110, 1010, 1910, 2810, and 3710 ms). Images were acquired with 1.3 mm³ isotropic resolution, and 144 sagittal slices were reconstructed.

MYC estimation was performed via multi-component fitting algorithms on derived maps. After acquisition, quantitative maps of T_1_), T_2_, and PD were computed and processed using the SyMRI software suite (SyntheticMR). These parametric maps were input into a multi-compartment Bloch equation–based signal modeling algorithm, which estimated the MVF for each voxel. The final MVF map reflects the proportion of voxel signal attributable to myelin, independently of confounding water or cellular content. This process yielded biologically interpretable and spatially resolved maps of MVF, which were then used in both ROI-level and voxel-wise analyses to assess myelin integrity across subjects [11,27–31].

#### Synthetic FLAIR and Double Inversion Recovery (DIR) Imaging

Synthetic contrast-weighted images, including FLAIR and Double Inversion Recovery (DIR), were also generated using the SyMRI software suite. Respective images were synthesized from the QALAS data by applying Bloch-equation–based signal modeling to voxel-wise T1, T2, and PD maps. The synthetic FLAIR images were configured with TR = 15,000 ms, TE = 90 ms, and T_I_ = 3,100 ms to suppress CSF. Synthetic DIR images were generated using TR = 15,000 ms, TE = 100 ms, with dual inversion times of TI₁ = 470 ms and TI₂ = 3,800 ms to simultaneously null white matter and CSF signals, thereby improving cortical gray matter contrast. Because all synthetic images are derived from the same acquisition, they are inherently co-registered, eliminating inter-scan motion artifacts, enabling pixel-wise comparison across contrasts.

#### FD Metric Derivation and Preprocessing

The FD map for each subject was computed voxel-wise as FD = (FLAIR – DIR) / FLAIR (Figure 1). This computation yields a normalized difference emphasizing the signal difference between FLAIR and DIR at each voxel. Prior to this calculation, several preprocessing steps were applied to ensure accurate correspondence between images and to stabilize the ratio. Each subject’s synthetic DIR image was rigidly coregistered to their synthetic FLAIR to ensure spatial alignment of corresponding voxels. Given that the synthetic images are intrinsically aligned, only minimal alignment adjustment was necessary. We then applied the same linear transformation to the MVF map to align it with the anatomical space.

**Figure 1.**
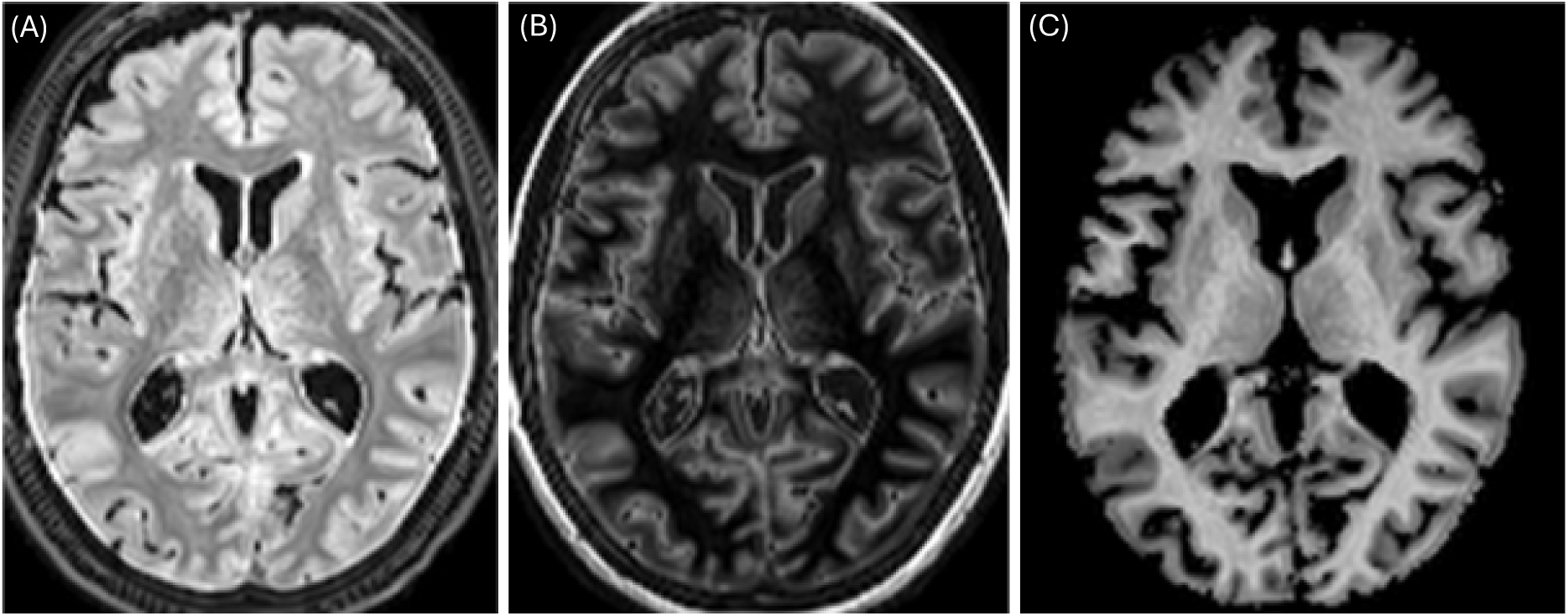
Illustration of the FLAIR–DIR/FLAIR ratio-based myelin mapping approach. Panel (A) shows the FLAIR image, which suppresses CSF and is sensitive to white matter signal. Panel (B) presents the DIR image, which suppresses both CSF and white matter, enhancing gray matter contrast. Panel (C) displays the resulting FD map computed as (FLAIR−DIR)/FLAIR.

A combined brain parenchyma mask (gray matter + white matter) was derived from each subject’s T1 segmentation. This mask was applied to both FLAIR and DIR images to exclude non-brain voxels and cerebrospinal fluid regions, focusing the FD calculation on brain tissue only. To avoid instabilities in the ratio when divided by very low FLAIR values, we excluded voxels with near-zero FLAIR intensity. Specifically, voxels with FLAIR signal below a small threshold were not evaluated, effectively setting FD undefined in areas outside the analysis mask. This typically removes only voxels in deep CSF or at brain edges where FLAIR is ∼0.

Noise and partial volume effects can lead to instances where DIR > FLAIR, resulting in a negative FD. Such negative FD values are not meaningful and likely indicate local intensity inversion due to noise or minor registration offsets. We set any FD < 0 to zero for all subjects, to ensure the FD metric reflects only positive or null differences. This was done consistently across the dataset.

Because FD is a relative contrast that is intensity-based and could have different overall ranges per subject, we linearly scaled each subject’s FD map to a common 0–1 range for voxel-wise statistical analyses. The min–max normalization was performed within that subject’s brain mask, so that the lowest FD in brain tissue became 0 and the highest became 1. This step mitigates inter-subject intensity drift and allows group-level, voxel-wise tests without bias from arbitrary intensity offsets.

#### Spatial Normalization

For group comparisons and voxel-wise analyses, individual FD and MVF maps were spatially normalized to a standard brain template (Montreal Neurological Institute; MNI space). Each subject’s T1-weighted image was non-linearly normalized to MNI space using SPM12’s diffeomorphic warping, and the resulting deformation fields were applied to the FD and MVF maps [32]. Images were resampled to the MNI template at ∼1.3 mm isotropic voxel size for statistical analysis. Prior to voxel-wise analyses, normalized FD and MVF maps were smoothed with an isotropic 4 mm full-width-at-half-maximum (FWHM) Gaussian kernel. Voxel-wise analyses were restricted to a combined gray- and white-matter mask incorporating a priori regions of interest relevant to AD pathophysiology, thereby focusing on brain parenchyma and excluding voxels in ventricles and outside the brain.

#### ROI Definition and Group Difference Analysis

We first examined ROI-based differences in FD and MVF between HC and MCI groups. ROI masks were defined a priori from standard brain atlases: for gray matter regions, we used the CONN atlas, and for white matter regions, we used the JHU white matter atlas [33,34]. In total, this yielded a comprehensive set of ROIs covering major cortical areas, limbic structures, and white-matter tracts. For each subject, we calculated the mean FD within each ROI and similarly extracted the mean MVF for each ROI. These ROI mean values were used to compare groups. Group comparisons (MCI vs HC) on ROI values were assessed using analysis of covariance (ANCOVA), with age included as a covariate. For each ROI, the ANCOVA tested the main effect of diagnosis (HC vs MCI) on the ROI’s FD or MVF, controlling for age. We considered results significant at p < 0.05 (uncorrected) at the ROI level; given the exploratory nature of this analysis, significant regions were later interpreted in context rather than strictly corrected for multiple comparisons due to the small sample.

#### ROI-Based Classification of MCI vs HC

To evaluate the discriminatory performance of FD and MVF, we conducted machine-learning classification analyses using ROI-based features. For each subject, feature vectors consisted of ROI values derived from the atlases described above.To reduce dimensionality, principal component analysis (PCA) was applied separately to each feature set (FD and MVF), and the minimum number of principal components explaining ≥85% of the variance was retained. This typically resulted in 5–8 components per feature set.

Classification was performed separately for FD-based and MVF-based features. Within each modality, three models were evaluated: *Model 1*: PCA components only. *Model 2*: PCA components plus age. *Model 3*: PCA components, age, and olfactory identification scores. The inclusion of age and odor ID in Models 2 and 3 allowed us to test if adding these subject-level variables improved classification. Classification was done using a logistic regression classifier trained to distinguish MCI vs HC. We employed a five-fold cross-validation scheme: the data were partitioned into 5 folds, the model was trained on 4 folds and tested on the held-out fold, and this was repeated so each subject was tested once. Model performance was aggregated across folds. We measured discriminative performance using the area under the Receiver Operating Characteristic curve (AUC). We report the pooled cross-validated AUC for each model. This procedure was run separately for the three ROI sets (All ROIs, GM-only, WM-only) to explore which tissue compartment yields the best classification. Additionally, to directly compare the utility of FD vs MVF, we ran an auxiliary set of classifications: FD-only features, MVF-only features, and combined FD+MVF features for Model 1 without covariates to see if combining both metrics improved discrimination.

#### Voxel-Wise Regression Analyses for FD and MVF vs Odor ID

We examined voxel-wise associations between FD/MVF and olfactory identification (ID) performance using DPABI [35]. The olfactory ID score was specified as the predictor of interest. *Model A (FD–ID, age-controlled)*: A voxel-wise regression was conducted with the ID score predicting FD at each voxel, controlling for age. Model B (FD–ID controlling for age and MVF): To test whether FD–ID associations were independent of local myelin estimates, MVF was included as a voxel-wise image covariate alongside age. A significant FD–ID effect in this model indicates variance in olfactory performance explained by FD beyond that attributable to MVF. For comparison, an age-controlled voxel-wise regression of MVF versus ID was also performed. Threshold for statistical maps was set at p < 0.005 (uncorrected), with cluster-extent thresholds set to k ≥ 40 voxels for FD models and k ≥ 10 voxels for MVF–ID. Given the modest sample size, these voxel-wise analyses were considered exploratory and were not corrected for multiple comparisons.

To assess spatial correspondence between FD and MVF, we computed voxel-wise Pearson correlations across subjects. For each voxel, correlation coefficients were calculated between FD and MVF values across participants, generating a map of shared inter-subject variance. Positive correlations indicated regional concordance between metrics, whereas weaker or negative correlations suggested divergence, informing interpretation of FD as a complementary measure.

## RESULTS

### Group differences in FD and MVF (ROI based analyses)

Two white matter ROIs demonstrated higher FD values in the HC compared to MCI (ANCOVA, p < 0.05, uncorrected, Figure 2). These included the splenium and body of the corpus callosum, indicating relatively lower FD values in posterior and mid-callosal fibers in MCI. Gray matter regions showing higher FD in HC comprised the left thalamus, bilateral hippocampus, left amygdala, left Frontal-orbital l cortex, and bilateral anterior olfactory nucleus (Figure 2A). Across these regions, the MCI group exhibited reduced FD, consistent with regional microstructural differences affecting both white and gray matter.

**Figure 2.**
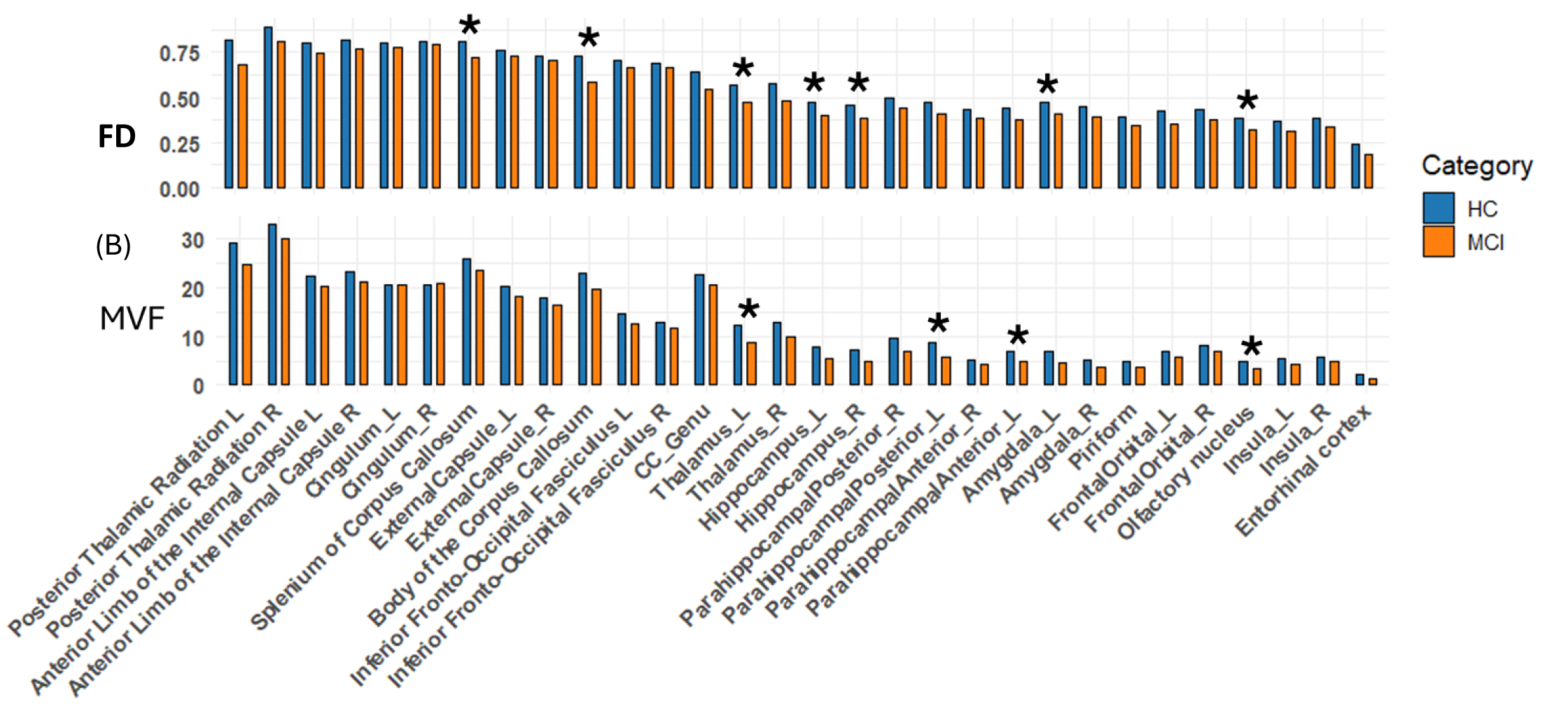
ROI-based group differences in MVF and FD between HC and MCI samples. (A) Mean FD computed as (FLAIR−DIR)/FLAIR by ROI, (B) Mean MVF for the same ROIs, plotted in the same left-to-right order to enable direct comparison across metrics. Age-adjusted ANCOVA (p < 0.05, uncorrected) showed higher FD in HC than MCI in the corpus callosum (splenium/body), left thalamus, left amygdala, bilateral hippocampus, bilateral anterior olfactory nucleus, and left orbitofrontal cortex. MVF was higher in HC in the left thalamus, left anterior and posterior parahippocampal gyrus, and bilateral anterior olfactory nucleus (p < 0.05, uncorrected).

In comparison, ROI-based MVF analyses revealed a more limited pattern of group differences. Higher MVF values in HC were observed in the left thalamus, left parahippocampal gyrus (anterior and posterior divisions), and bilateral anterior olfactory nucleus (p < 0.05, uncorrected) (Figure 1B). Although there was partial overlap with FD findings, MVF did not demonstrate group differences in the hippocampi or corpus callosum under the same statistical threshold.

### ROI-Based Classification of MCI vs HC

FD-based classification demonstrated moderate-to-good discrimination across ROI groups (Table 2). For all ROIs, AUCs were 0.757 for Model-1, 0.691 for Model-2, and 0.704 for Model-3. For gray-matter ROIs, AUCs were 0.779 (Model-1), 0.761 (Model-2), and 0.757 (Model-3). For white-matter ROIs, AUCs were 0.739 (Model-1), 0.746 (Model-2), and 0.713 (Model-3).

**Table 2.**
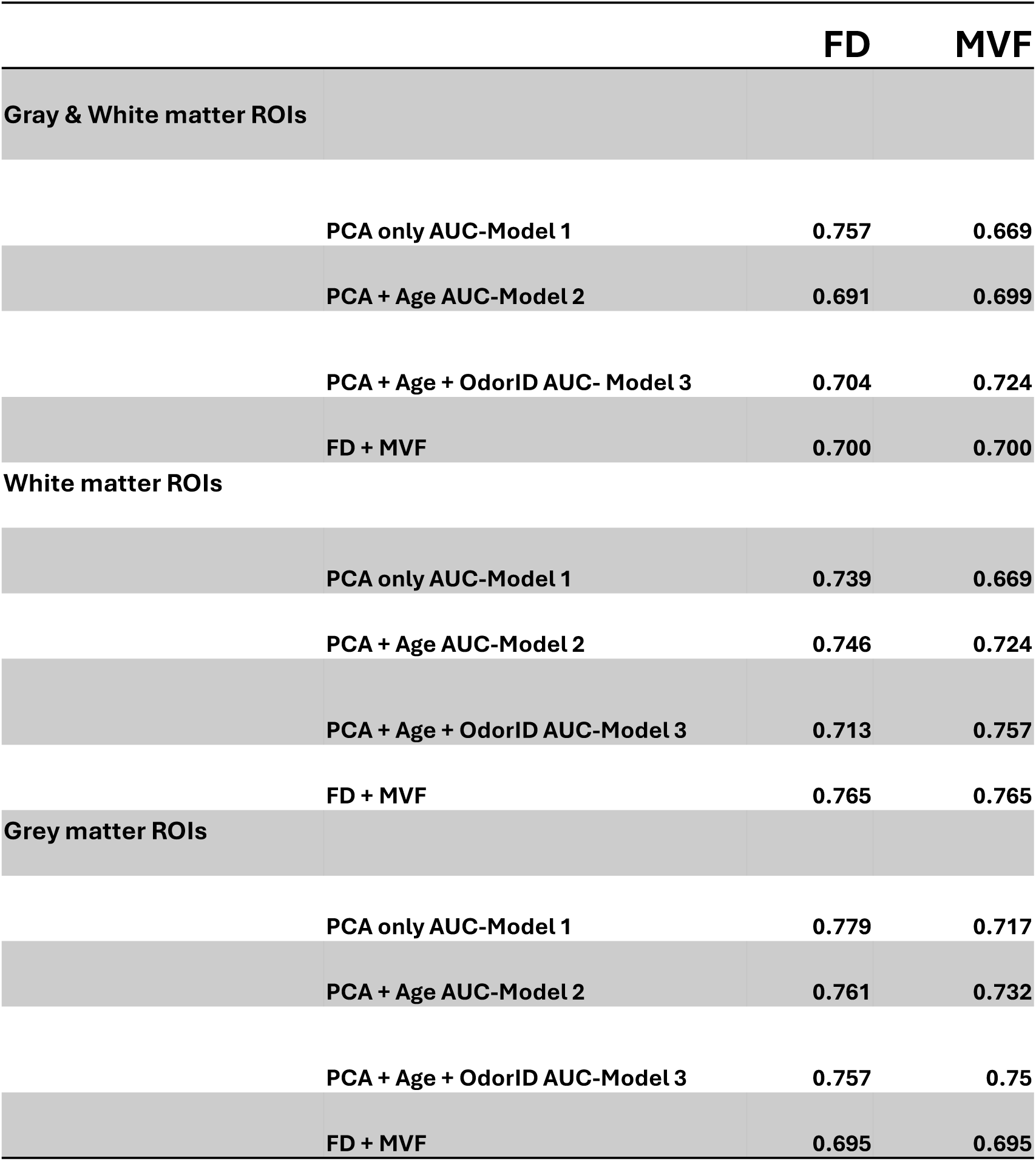
ROI-based classification of MCI vs HC using FD and MVF. Cross-validated AUCs are shown for all, gray-matter, and white-matter ROI sets. Model-1 uses PCA-reduced ROI features, Model-2 adds age, and Model-3 adds age + odor identification (ID).

For FD, peak performance was observed in gray matter ROIs under Model-1 (PCA features only; AUC = 0.779). Inclusion of age (Model-2) or age plus odor identification (Model-3) did not improve FD-based performance and slightly reduced AUC in the all-ROI model, consistent with FD features already capturing covariate-related variance relevant to group separation and with potential redundancy introduced by additional predictors in a relatively modest sample.

In contrast, MVF-based classification improved with covariate inclusion. For all ROIs, AUCs increased from 0.669 (Model-1) to 0.724 (Model-3). Similar improvements were observed in gray matter ROIs (0.717 to 0.750) and white matter ROIs (0.669 to 0.757), indicating that adding odor identification enhanced MVF’s discriminative performance.

Direct comparisons showed numerically higher baseline AUCs for FD-only than MVF-only models across ROI sets in this sample. For example, using all ROIs without covariates, FD achieved AUC = 0.757 versus MVF = 0.669; in gray-matter ROIs, FD achieved 0.779 versus MVF = 0.717. Combining FD and MVF features did not yield meaningful gains, producing intermediate AUCs (∼0.70–0.76) that were comparable to or slightly below FD alone. Collectively, these results indicate that ROI-level FD and MVF patterns captured overlapping but not identical information, with FD showing numerically higher baseline discrimination in this dataset, supporting FD as a potentially sensitive biomarker of early disease-related alteration.

### Voxel-Wise Associations with Olfactory Identification for FD and MVF

We next examined voxel-wise relationships between FD, MVF, and olfactory identification performance while controlling for age (Figure 3). MVF–Odor ID (age-adjusted) showed significant positive associations in the insula and the genu of the corpus callosum (p < 0.005, uncorrected; k ≥ 10 voxels; Figure 3 A). This pattern suggests that higher myelin-related signal is linked to better odor identification, with effects spanning both a cortical region implicated in chemosensory/salience integration, i.e. insula, and the genu callosum, a frontal interhemispheric white-matter pathway.

**Figure 3.**
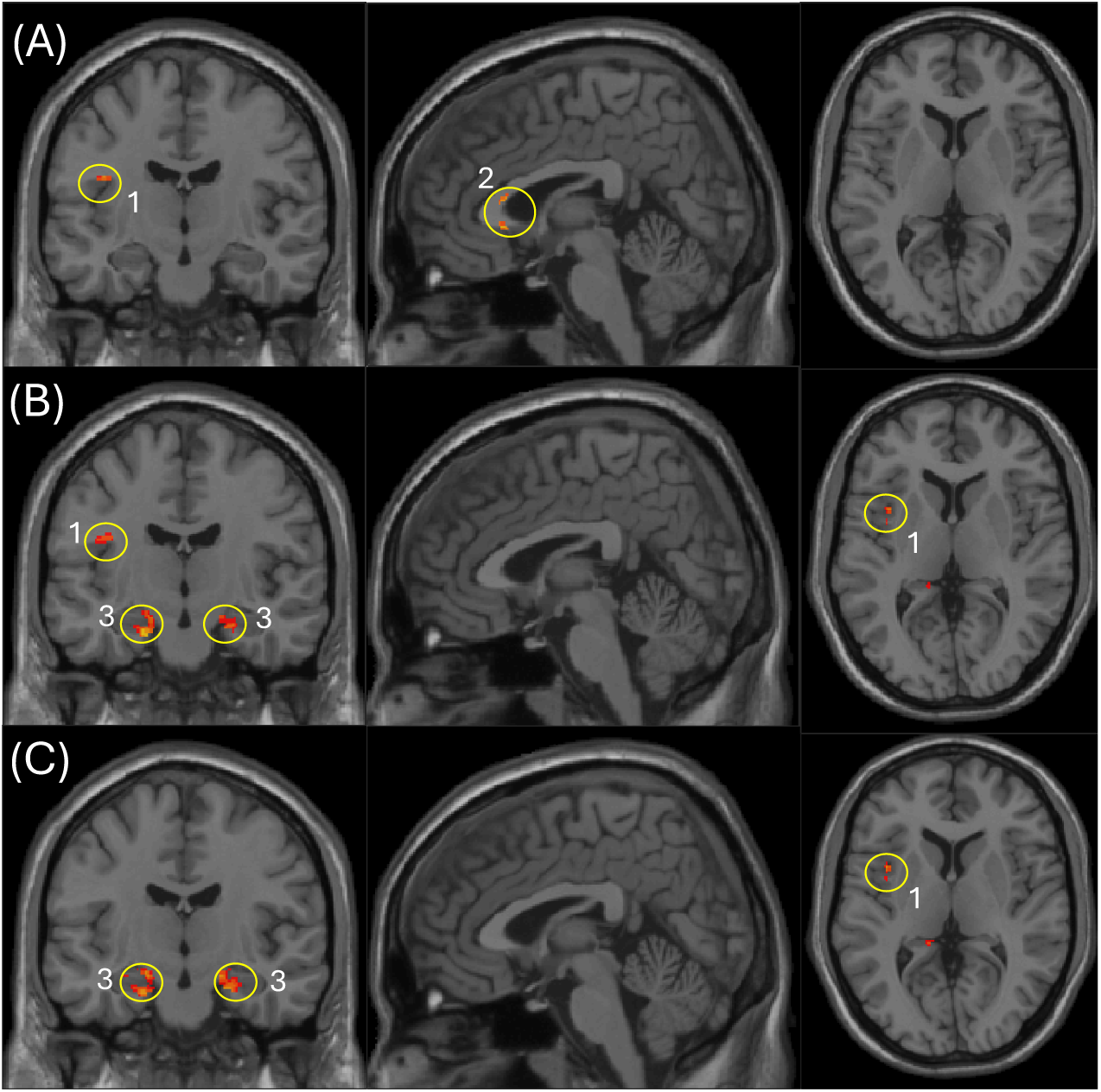
Voxel-wise regression and cross-modal analyses. Panels A–C show age-adjusted voxel-wise associations with odor identification (ID). (A) Positive MVF–ID associations (k ≥ 10 voxels) in the insula and genu of the corpus callosum. (B) Positive FD–ID associations (k ≥ 40 voxels) in the insula and hippocampus/parahippocampal regions. (C) Positive FD–ID associations after additionally including voxel-wise MVF as an image covariate (age + MVF adjusted), showing clusters in the insula and hippocampus/parahippocampal regions. (1) Insula, (2) Genu of corpus callosum, (3) Hippocampus and parahippocampal regions.

**I**n contrast, FD–Odor ID relationship (age-adjusted) demonstrated significant positive clusters in the insula and hippocampus (p < 0.005, uncorrected; k ≥ 40 voxels; Figure 3 B). The hippocampal involvement aligns with medial temporal contributions to odor–memory, while the insula finding supports a role for higher-order sensory integration in individual differences during odor identification [36,37].

To test whether FD’s association with odor ID was independent of local myelin estimates, we repeated the FD model including MVF as a voxel-wise image covariate (Figure 3 C). In this analysis, FD remained positively associated with odor ID in the hippocampus and insula (p < 0.005, uncorrected, using the same inference framework as above). This indicated that FD captures variance related to olfactory performance in medial temporal and insular regions that are not fully explained by MVF, consistent with FD reflecting complementary tissue-property sensitivity beyond myelin. Overall, these findings highlight partially overlapping but non-identical FD and MVF correlates of olfactory identification, with FD showing prominent associations in hippocampal–insular circuitry and MVF showing additional associations in callosal white matter.

### Cross-Modal Correspondence between FD and MVF

We assessed how closely FD maps resembled MVF maps across subjects using voxel-wise Pearson correlation analysis. Correlation coefficients were computed at each voxel across participants, generating a map of shared inter-subject variance. For this cross-modal correspondence analysis, the resulting map was thresholded at p < 0.0001 (uncorrected) with a minimum cluster extent of 100 voxels. We found predominantly positive correlations between FD and MVF across wide areas of the brain, confirming that FD is broadly related to myelin content. Strong positive correspondence was observed in regions such as the hippocampus, insula, thalamus, superior corona radiata, and posterior limb of the internal capsule (Figure 4). Subjects with higher MVF in the thalamus or internal capsule also tended to have higher FD in those same regions, consistent with both metrics increasing with greater myelin-related structural integrity. This spatial convergence suggests that FD partially indexes myelin content, strengthening its validity as a myelin-sensitive measure. However, the correlation was not uniform across the brain; some regions showed only modest or no correlation, indicating that FD is not a trivial surrogate for MVF. Notably, cortical gray-matter regions often showed lower FD–MVF correlation, likely reflecting the relatively limited dynamic range of MVF in cortex, whereas FD may capture additional microstructural variation through its DIR-derived signal properties.

**Figure 4.**
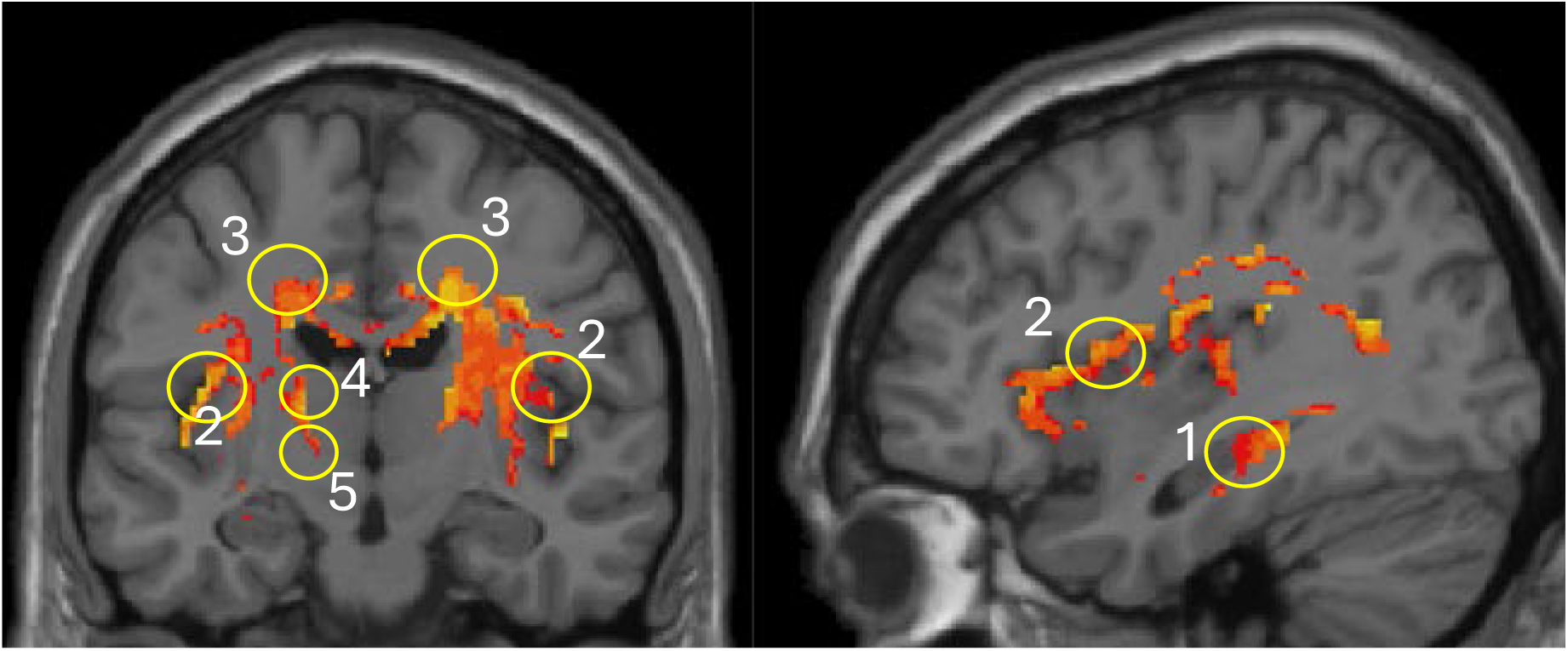
Positive voxel-wise correspondence between FD and MVF. (1) Hippocampus and parahippocampal regions, (2) Insula, (3) Superior corona radiata, (4) Thalamus, (5) Posterior limb of internal capsule.

## DISCUSSION

Our findings demonstrated that the synthetic FLAIR–DIR–derived metric FD sensitively captures MCI-related brain alterations, including the core behavioral phenotype of olfactory dysfunction. FD identified abnormalities in interconnected white matter and limbic regions—including the corpus callosum, hippocampus, amygdala, orbitofrontal cortex, thalamus, and olfactory nucleus—closely aligning with established models of AD propagation that emphasize early involvement of olfactory–limbic circuits and subsequent network-level spread [38,39].

We observed lower FD values in MCI compared with HC. Because FD reflects differences in inversion-recovery signal behavior between FLAIR and DIR contrasts, which are governed by T1 recovery and T2 decay processes, this finding suggests that early microstructural alterations may reduce the differential signal response between these sequences. Subtle changes in myelin organization, lipid composition, or tissue water balance may modify relaxation behavior in a manner that brings FLAIR and DIR signals closer together, resulting in lower FD values. At the same time, higher FD values in MCI are also biologically plausible, as more pronounced microstructural disruption—such as demyelination or increased extracellular water—could amplify differences between these contrasts. Consequently, FD may vary in either direction depending on the dominant tissue changes and disease stage, potentially reflecting the heterogeneous microstructural processes (Figure 5).

**Figure 5.**
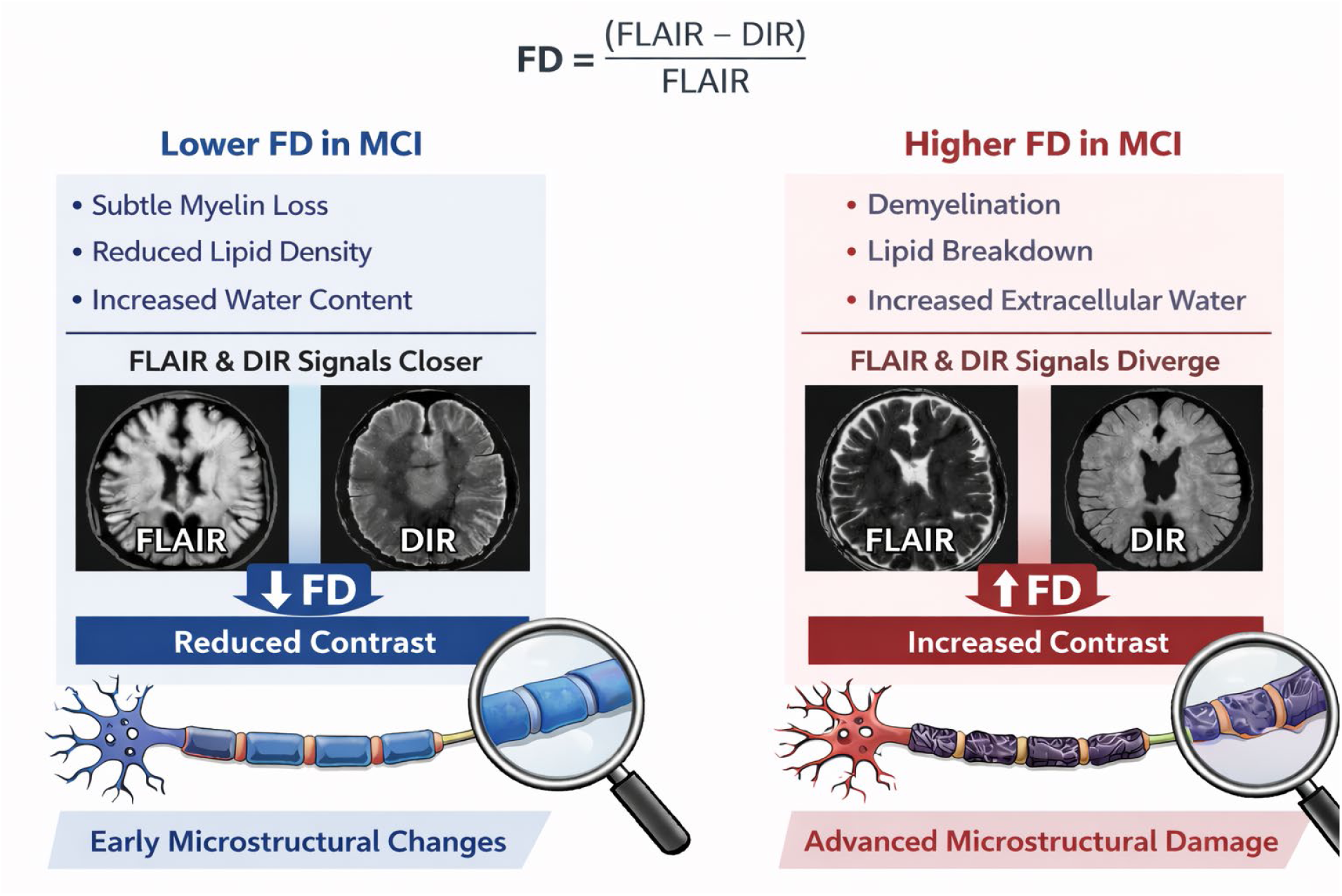
Conceptual model for FD in MCI. Changes in tissue microstructure may differentially affect FLAIR and DIR signals, producing either decreased or increased FD values depending on the nature and stage of the underlying myeline alterations.

Importantly, FD performed equally well to that of MVF in distinguishing individuals with MCI from HC controls. It demonstrated robust associations with olfactory identification performance, including in regions where MVF showed a more focal pattern of associations. In voxel-wise analyses, FD–odor identification associations were observed prominently in the insula and hippocampus/parahippocampal areas, whereas MVF showed more focal associations, for example, in the insula and genu of the corpus callosum. Notably, FD–odor associations in medial temporal and insular regions persisted after adjusting for voxel-wise MVF. Therefore, FD seems to capture complementary microstructural alterations potentially related to olfactory performance that are not fully captured by conventional myelin-sensitive measures such as MVF.

Higher FD–odor identification correlation within medial temporal regions is consistent with its role in both memory and olfactory processing. The insular association supports FDs’ involvement in higher-order sensory integration and salience-related circuitry during odor identification. Likewise, MVF demonstrated positive associations with odor identification; however, these were comparatively more focal, with clusters observed in the insula and genu of the corpus callosum. The shared insular involvement suggests that both FD and MVF capture olfaction-relevant tissue properties in this region, whereas the MVF association in the anterior corpus callosum is consistent with the interpretation that greater callosal myelin content may support efficient interhemispheric and frontal network communication relevant to olfactory performance [36,37,40,41].

Notably, MVF did not demonstrate significant voxel-level associations within hippocampal or other gray-matter regions at the same statistical threshold. In contrast, FD detected a hippocampal relationship not observed with MVF, suggesting sensitivity to olfaction-related microstructural variation within medial temporal circuitry that was not fully captured by MVF. Together, these findings indicated that while FD shares myelin-related sensitivity with MVF in white matter, it provides additional gray-matter–relevant information that may reflect early network vulnerability in MCI.

Viewed through a network-based perspective of AD progression, these findings position FD as a promising imaging marker capable of indexing potential microstructural disruption along vulnerable olfactory–limbic pathways. By linking such structural alterations to behavioral impairments, FD may enhance sensitivity to early disease-related changes and contribute to biomarker development aimed at identifying individuals at risk along the AD continuum [42].

### FD as a Myelin/Lipid-Sensitive Marker

Because FD is derived from FLAIR and DIR contrasts, it emphasizes tissue compartments in which FLAIR retains signal while DIR suppresses normal-appearing white matter. Higher FD values therefore may reflect greater contributions from myelin- and lipid-rich tissue components [9,10,13,14,16,17,43]. However, FD is not specific to myelin and may also be influenced by inflammation, gliosis, water content, or partial volume effects [44]. Accordingly, FD should be interpreted as a semi-quantitative, myelin-/lipid-sensitive contrast rather than a direct measure of myelin content.

The positive correspondence between FD and MVF in major white-matter pathways, however, supports its sensitivity to myelin-related variance. This finding is consistent with prior work showing that combinations of conventional MRI contrasts—such as the T1w/T2w ratio—can generate myelin-sensitive maps [2,13,26], and with reports of strong associations between T1w/T2w and SyMRI-derived MVF in demyelinating disease cohorts [1]. At the same time, the biological interpretation of ratio-based metrics is context dependent, and discrepancies with histology and technical confounds have been described [13,45], underscoring the need for cautious inference.

Notably, FD demonstrated group differences within hippocampal and cortical regions that were less prominent for MVF, suggesting sensitivity to gray-matter properties not fully captured by MVF. Ratio-based approaches have previously been used to characterize intracortical microstructural variation [2,46], supporting the plausibility of detecting depth- and region-dependent differences within gray matter.

Critically, the FD–olfactory identification association in the hippocampus/parahippocampal regions and insula persisted after adjustment for MVF, indicating that FD captures variance beyond local MVF estimates. Medial temporal regions are characterized by high synaptic density, lipid-rich neuronal membranes, and metabolically active circuits that are vulnerable to activity-dependent stress and trans-synaptic propagation of Alzheimer’s disease pathology. Early AD is associated with the disruption of axonal membranes, myelinated fibers, and lipid homeostasis within limbic regions, potentially preceding overt neuronal loss. FD may therefore index membrane-and lipid-related microstructural alterations contributing to early network dysfunction affecting olfaction [3–5,16,17,43].

Consistent with this framework, better olfactory identification was associated with higher FD in medial temporal regions such as the hippocampal/parahippocampal and insular regions. Notably, these relationships remained evident even after additional adjustments for voxel-wise MVF. In comparison, MVF showed positive voxel-wise associations with odor identification that were more focal, with effects observed in the insula and genu of the corpus callosum. Overall, FD and MVF appear to share sensitivity to common contrast mechanisms in white matter, while FD shows additional olfaction-relevant associations in medial temporal circuitry and insular cortex that are not fully accounted for by MVF. This pattern suggests that FD is sensitive to tissue properties that modulate effective T1/T2 provides complementary, functionally relevant information and may serve as a myelin-/lipid-sensitive marker of early medial temporal vulnerability.

Beyond MVF and T1w/T2w-style ratios, several other MRI approaches have been used to probe myelin, including magnetization transfer–based methods such as MTR, MTsat, and ihMT, myelin water imaging, susceptibility-based methods such as quantitative susceptibility mapping, and diffusion-informed composite measures such as the g-ratio [47–50]. These techniques differ in the tissue compartment they emphasize, with some being more sensitive to macromolecular or bound proton content, others to water trapped between myelin bilayers, and others to susceptibility or axon–myelin relationships [51,52]. Accordingly, they provide complementary rather than interchangeable information, and their relative performance depends on the biological question, acquisition constraints, and processing complexity. Within this broader framework, FD may be best viewed not as a replacement for dedicated quantitative myelin imaging, but as a pragmatic semi-quantitative marker derived from clinically accessible synthetic inversion-recovery contrasts that can complement MVF while remaining feasible in settings where more specialized myelin protocols are unavailable [9,47].

### Clinical Relevance to Olfactory Association

The observed associations between FD in hippocampal and callosal regions and odor identification are clinically relevant in the context of Alzheimer’s disease risk. Odor identification impairment is a well-established early feature across the AD spectrum and has been shown to predict incident MCI and subsequent progression to AD dementia [25,53]. Neuroimaging studies further demonstrate that poorer odor identification is linked to structural vulnerability in olfactory–limbic circuitry, including the olfactory cortex, amygdala, entorhinal cortex, hippocampus, and insula [21,54]. Postmortem evidence also supports olfactory pathway involvement in AD, including substantial loss of myelinated axons in the olfactory tract [24]. Consistent with this literature, the persistence of FD–odor identification associations in the hippocampus after adjustment for voxel-wise MVF suggests that FD may capture tissue-property variance relevant to olfactory–memory circuit integrity beyond local quantitative MVF estimates. In contrast, the more limited MVF–odor identification effects observed in callosal regions may indicate that olfactory performance relates not only to bulk white-matter myelin content but also to gray-matter and limbic microstructural integrity that is variably captured by MVF [21,54–57]. Together, these findings support the potential utility of FD as a pragmatic imaging-derived marker for characterizing olfactory-network vulnerability in MCI, with implications for risk stratification that merit validation in larger and longitudinal cohorts [20,25].

### FD and MVF: Comparison and Complementarity

FD-based ROI patterns showed numerically higher baseline classification performance than MVF in this cohort, while the two measures appeared to provide overlapping and complementary information. This difference may reflect both measurement characteristics and tissue sensitivity. Although MVF provides a quantitative estimate of myelin-related volume, SyMRI-derived compartment modeling can be affected by tissue-boundary partial-volume effects, and quantifying myelin-related signal in cortex is intrinsically challenging because gray matter contains relatively low myelin content and thinner laminar structure [10,58]. In contrast, FD is derived from inversion-recovery contrasts (FLAIR and DIR) that yield strong tissue contrast through CSF and white-matter suppression mechanisms, potentially increasing sensitivity to subtle intensity differences in regions with complex tissue composition [10]. FD also reflects combined contributions from white and gray matter, whereas MVF contrast and dynamic range are typically most pronounced in white matter, which may reduce sensitivity to gray-matter–predominant alterations. Consistent with this interpretation, adding MVF to FD did not improve cross-validated performance, suggesting substantial overlaps in discriminative information at the ROI level. At the same time, the positive FD–MVF correspondence observed in several regions supports that FD captures myelin-related variance, aligning with prior work showing that combining conventional MRI contrasts can yield myelin-sensitive maps such as T1w/T2w.

The anatomical distribution of FD findings is consistent with reported vulnerability of olfactory–limbic circuitry in prodromal AD. Reduced FD in the corpus callosum (splenium/body) is compatible with callosal involvement in AD-spectrum conditions and has been linked to interhemispheric disconnection-related mechanisms [40,41]. FD differences in the hippocampus and parahippocampal regions—key nodes of medial temporal networks implicated early in tau-related neurodegeneration—support sensitivity to microstructural alterations in memory circuitry [59–61]. The persistence of FD–odor identification associations in hippocampal regions after controlling for voxel-wise MVF further suggests that FD captures variance related to olfactory performance not fully explained by local MVF estimates. Thalamic FD and MVF effects may reflect involvement of thalamo-cortical and limbic relay systems, which have also been reported to show early structural change in MCI/AD [62,63]. Finally, FD differences in the orbitofrontal cortex and anterior olfactory nucleus are consistent with involvement of higher-order olfactory and olfactory-network hubs implicated in AD-related olfactory dysfunction [64–66]. Together, these patterns indicate that FD highlights a set of white- and gray-matter regions relevant to olfaction and memory and may provide a practical imaging readout of integrity within this network.

Several limitations should be considered. First, the sample size (n=33) limits statistical power and generalizability. ROI and voxel-wise results were evaluated using uncorrected thresholds in an exploratory framework; replication in larger independent samples will be necessary to establish robustness. Second, FD is a semi-quantitative intensity-derived metric and is dependent on the stability of synthetic FLAIR and DIR generation; scanner- and site-related variability could influence FD scaling, although within-subject normalization mitigates some effects. Third, while FD showed partial convergence with MVF, we did not perform histopathologic validation, and FD may reflect additional tissue properties, such as water content or inflammatory changes beyond myelin/lipid content. Fourth, MVF derived from synthetic MRI, while supported by prior validation studies, may underrepresent gray-matter myelin and remain susceptible to partial-volume effects, which could reduce sensitivity in cortical/medial temporal regions. Finally, odor identification is influenced by non-neurological factors such as sinonasal disease, which may introduce noise into brain–behavior associations.

Future work should evaluate FD longitudinally to determine whether FD trajectories track cognitive decline or conversion from MCI to dementia and whether baseline FD in olfactory–limbic regions predict outcomes including integrating FD with complementary modalities such as DTI microstructure and amyloid/tau PET when available and clarifying whether FD aligns more closely with white-matter integrity, molecular pathology, or both. Additional technical work is also warranted to quantify test–retest reliability, optimize FD computation and normalization, and assess cross-scanner harmonization. Finally, validating FD in additional cohorts and disease contexts characterized by myelin-related changes would help define its specificity and clinical utility.

These findings indicated that individuals with preserved or higher FD in these regions tended to have higher odor identification performance. The hippocampal finding aligns with the role of the hippocampus and associated medial temporal lobe structures in both memory and olfactory processing. The callosal finding suggests that white-matter integrity in frontal interhemispheric connections may also relate to olfactory function. For comparison, MVF showed positive associations with odor ID as well, but the pattern was more limited. A cluster of positive MVF–ID correlation was observed in the genu of the corpus callosum, overlapping with where FD showed an effect. This suggests that higher myelin content in the anterior corpus callosum is linked to better odor identification, as the olfactory and orbitofrontal regions communicate via these fibers. However, MVF did not show significant voxel-level associations in the hippocampal or other gray-matter regions at the same threshold – the effects were comparatively sparse, mostly confined to some white-matter areas like the genu. The contrast between these results suggests that FD may capture partially distinct microstructural features from MVF to olfaction-related microstructural differences in certain regions. FD detected a relationship in the hippocampus that MVF did not, potentially because FD captures additional tissue property differences in hippocampal circuitry.

## Conclusion

FD represents a novel MRI-derived metric with potential utility for detecting subtle brain alterations in MCI related to tissue integrity and myelin-sensitive properties. In this study, FD differentiated individuals with MCI from HCs across several AD-relevant brain regions and showed associations with olfactory identification performance that were partially overlapping with, but not identical to, those observed with MVF. Because FD reflects differences in inversion-recovery signal behavior between FLAIR and DIR contrasts, which are governed by T1 recovery and T2 decay processes, it is potentially sensitive to microstructural variations affecting myelin integrity, lipid composition, and tissue water content [67]. These findings suggest that FD captures early alterations in tissue relaxation behavior and may provide a practical, scalable marker of heterogeneous microstructural processes occurring during the early stages of AD.

In our analysis, FD showed partial spatial overlap with MVF—supporting its sensitivity to myelin-related characteristics. However, it also captured additional variance, particularly within gray matter regions. This pattern suggests that FD may reflect complementary microstructural information not fully represented by conventional quantitative myelin measures such as MVF.

Given that FD can be derived from routinely acquired FLAIR and DIR sequences, it may offer a practical and accessible addition to the neuroimaging toolkit for probing brain health. Within the context of AD research, FD could serve as a surrogate marker to enhance early detection efforts and support risk stratification in individuals along the cognitive decline continuum.

