## supplementary material for "A DERIVED RELAXATION CONTRAST FROM SYNTHETIC MRI FOR DETECTING NETWORK MICROSTRUCTURAL VULNERABILITY"

### SEMI-QUANTITATIVE CONTRAST FOR MYELIN/LIPID QUANTIFICATION USING SYNTHETIC FLAIR AND DIR IN MCI

**SUPPLEMENTARY MATERIAL**

**Running title:** Myelin changes in MCI

**Author affiliations:**

[1]Department of Radiology, Penn State University College of Medicine, Hershey, PA, USA

[2]Department of Neurology, Penn State University College of Medicine, Hershey, PA, USA

[3]Department of Neurosurgery, Penn State University College of Medicine, Hershey, PA, USA

#### FLAIR signal model

Using standard approximation:

$$S_{\text{FLAIR}} \approx M_0\left(1 - 2e^{-TI_F/T1} + e^{-TR/T1}\right)e^{-TE_F/T2}$$

let

$$A_F(T1) = \left(1 - 2e^{-TI_F/T1} + e^{-TR/T2}\right)$$

then

$$S_{\text{FLAIR}} \approx M_0\, A_F(T1)\, e^{-TE_F/T2}.$$

#### DIR signal model (double inversion)

A common simplified DIR approximation is:

$$S_{\text{DIR}} \approx M_0\left(1 - 2e^{-TI_1/T1} + 2e^{-TI_2/T1} - e^{-TR/T1}\right)e^{-TE_D/T2}$$

Define:

$$A_D(T_1) = \left(1 - 2e^{-TI_1/T1} + 2e^{-TI_2/T1} - e^{-TR/T1}\right)$$

then

$$S_{\text{DIR}} \approx M_0\, A_D(T1)\, e^{-TE_D/T2}.$$

Similarly, DF definition:

$$FD = \frac{\text{FLAIR} - \text{DIR}}{\text{FLAIR}} = 1 - \frac{\text{DIR}}{\text{FLAIR}} = 1 - \frac{S_{\text{DIR}}}{S_{\text{FLAIR}}}.$$

Substitute respective signal models:

$$FD = 1 - \frac{M_0\, A_D(T1)\, e^{-TE_D/T2}}{M_0\, A_F(T1)\, e^{-TE_F/T2}}$$

$$FD = 1 - \frac{A_D(T1)}{A_F(T1)}\; e^{-(TE_D - TE_F)/T2}.$$

The closed-form DF expression is:

$$FD = 1 - \frac{\left(1 - 2e^{-TI_1/T1} + 2e^{-TI_2/T1} - e^{-TR/T1}\right)}{\left(1 - 2e^{-TI_F/T1} + e^{-TR/T1}\right)}\; e^{-(TE_D - TE_F)/T2}$$

#### Conceptual interpretation of FD

The FD metric is essentially measuring differences in inversion-recovery contrast between FLAIR and DIR which are governed by:

- tissue T1 recovery
- T2 decay
- myelin / lipid microstructure

FD therefore behaves like a derived relaxation contrast reflecting network microstructural integrity

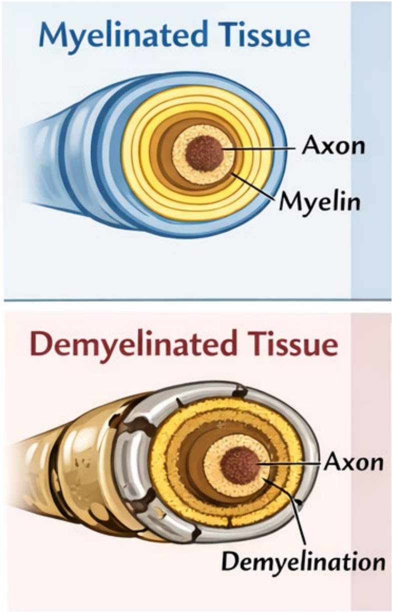


**Figure S1**. Conceptual illustration of low and high FD values in MCI. Variations in tissue microstructure may differentially affect FLAIR and DIR signals, leading to either reduced or increased FD values depending on the nature and stage of underlying microstructural alterations.

.

#### Mathematical Meaning T1/T2 ratio

T1= longitudinal (spin-lattice) relaxation time

T2= transverse (spin-spin) relaxation time

Both are measured in milliseconds, so the units cancel, making the ratio dimensionless.

| Tissue | T1 (ms) | T2 (ms) | T1/T2 |
|---|---|---|---|
| White matter | ~800 | ~80 | ~10 |
| Gray matter | ~1300 | ~100 | ~13 |
| CSF | ~4000 | ~2000 | ~2 |

Different tissues have different T1/T2 ratios, which contributes to MRI contrast. It is sensitive to:

- Myelin disruption
- lipid changes
- microstructural alterations

#### The relationship between FD and T1/T2 Ratio

| Tissue Property | Effect |
|---|---|
| Myelin | Short T1, short T2 |
| Lipid content | Alters both |
| Water content | Long T1, long T2 |

Microstructural disruption changes both relaxation times

Therefore:

$$DF = f(T1, T2)$$

which indirectly relates to the T1/T2 ratio, a known myelin-sensitive contrast. Therefore, FD can be considered sensitive to tissue properties that modulate effective $T1/T2$—including myelin- and lipid-related microstructural alterations—while remaining scalable because it is computed from routinely generated synthetic contrasts.
